# Elastin-derived peptides favor type 2 innate lymphoid cells in COPD

**DOI:** 10.1101/2023.09.13.557567

**Authors:** Sarah Lahire, Caroline Fichel, Océane Rubaszewski, Cédric Lerévérend, Sandra Audonnet, Vincent Visneux, Jeanne-Marie Perotin, Gaëtan Deslée, Sébastien Le Jan, Stéphane Potteaux, Richard Le Naour, Arnaud Pommier

## Abstract

Chronic obstructive pulmonary disease (COPD) is a condition characterized by chronic airway inflammation and obstruction, primarily caused by tobacco smoking. Although the involvement of immune cells in COPD pathogenesis is well established, the contribution of innate lymphoid cells (ILC) remains poorly understood. ILC are a type of innate immune cells that participate in tissue remodeling processes, but their specific role in COPD has not been fully elucidated. During COPD, the breakdown of pulmonary elastin generates elastin peptides that elicit biological activities on immune cells. This study aimed to investigate the presence of ILC in COPD patients and examine the impact of elastin peptides on their functionality.

Our findings revealed an elevated proportion of ILC2 in the peripheral blood of COPD patients, and a general activation of ILC as indicated by an increase in their cytokine secretion capacity. Notably, our study demonstrated that serum from COPD patients promotes ILC2 phenotype, likely due to the elevated concentration of IL-5, a cytokine known to favor ILC2 activation. Furthermore, we uncovered that this increase in IL-5 secretion is partially attributed to its secretion by macrophages upon stimulation by elastin peptides, suggesting an indirect role of elastin peptides on ILC in COPD.

These findings shed light on the involvement of ILC in COPD and provide insights into the potential interplay between elastin breakdown, immune cells, and disease progression. Further understanding of the mechanisms underlying ILC activation and their interaction with elastin peptides could contribute to the development of novel therapeutic strategies for COPD management.

## INTRODUCTION

Chronic obstructive pulmonary disease (COPD) is a chronic inflammatory disease of the airways that is mainly caused by tabacco smoking. COPD is very common. Indeed, it is now the third leading cause of death in the world [1]. COPD is defined by two factors: 1. A thickening of the bronchus wall and mucus hypersecretion that lead to a progressive narrowing and permanent obstruction of the airways and 2. An emphysema caused by the destruction of pulmonary alveoli that leads to shortness of breath [2, 3].

COPD features chronic inflammation of the lungs with a progressive infiltration of immune cells, both innate and adaptive, into the lung parenchyma. Whereas neutrophils and macrophages are key effector cells involved in COPD pathogenesis, recent studies have emphasized on the role of the persistent infiltration of other immune cells such as NK cells, NKT cells, and T lymphocytes in the pulmonary inflammation process and emphysema development [4–7]. The resulting chronic lung inflammation magnitude correlates with disease severity in COPD patients. Massive protease synthesis by lung-infiltrating inflammatory cells [4] results in pulmonary elastin breakdown that leads to emphysema and the generation of soluble elastin peptides (EP) [6, 8, 9]. EP exert their biological activities via their interaction with the elastin receptor complex (ERC) and galectin-3 [4]. We have shown both in a mouse model of EP-induced emphysema and in COPD patients, that EP participate in the pathophysiology of COPD. Indeed, EP modulate the reactivity of neutrophils [9] and macrophages [10], and influence the polarization of T cells [4, 6]. The potential effect of EP on innate lymphoid cells (ILC) in the context of COPD remains to be elucidated.

ILC are innate immune cells that play a central role in tissue remodelling associated with many pathologies. These cells are divided into three subgroups, ILC1, ILC2 and ILC3, that are characterized by their functions associated respectively with Th1-type, Th2-type and Th17-type helper T lymphocyte responses [11]. While ILC3 is the most represented population in the lung parenchyma, ILC1 and NKp46^-^ ILC3 cells accumulate in the lung of COPD patients [12, 13]. In a recent study, an increase in IL-33 serum levels associated with an increase in the proportion of ILC2 in the blood of COPD patients was shown [14]. Exposure of mice to cigarette smoke lead to an increase of IL-13, IL-33 and ILC2 in the lung, however ILC2 deficiency lead to impaired lung functions and lung hyperresponsiveness [15]. These results show that the contribution of ILC during COPD is yet to be completely elucidated. Here, we investigated the orientation of ILC in COPD patients and the impact of EP on them.

## MATERIALS AND METHODS

### Patients’ characteristics (Table 1)

This single-center study (NCT02924818) was conducted in the service of Pulmonary Medicine at the University Hospital of Reims, under the approval of an ethics committee in human biomedical research (CPP Dijon EST I, No. 2016-A00242-49). All subjects gave their informed and written consent prior to inclusion in the study. COPD patients were enrolled in the study based on clinical assessments with a forced expiratory volume in 1−s (FEV_1_)/forced vital capacity (FVC) < 0.7 after bronchodilation. At inclusion, all patients were stable for at least 4 weeks. Patients with asthma, allergic disease, tuberculosis, neoplasia, or other chronic respiratory diseases were excluded. Control subjects were recruited from the French Blood Institution and did not present any acute or chronic respiratory disease.

**Table 1:**
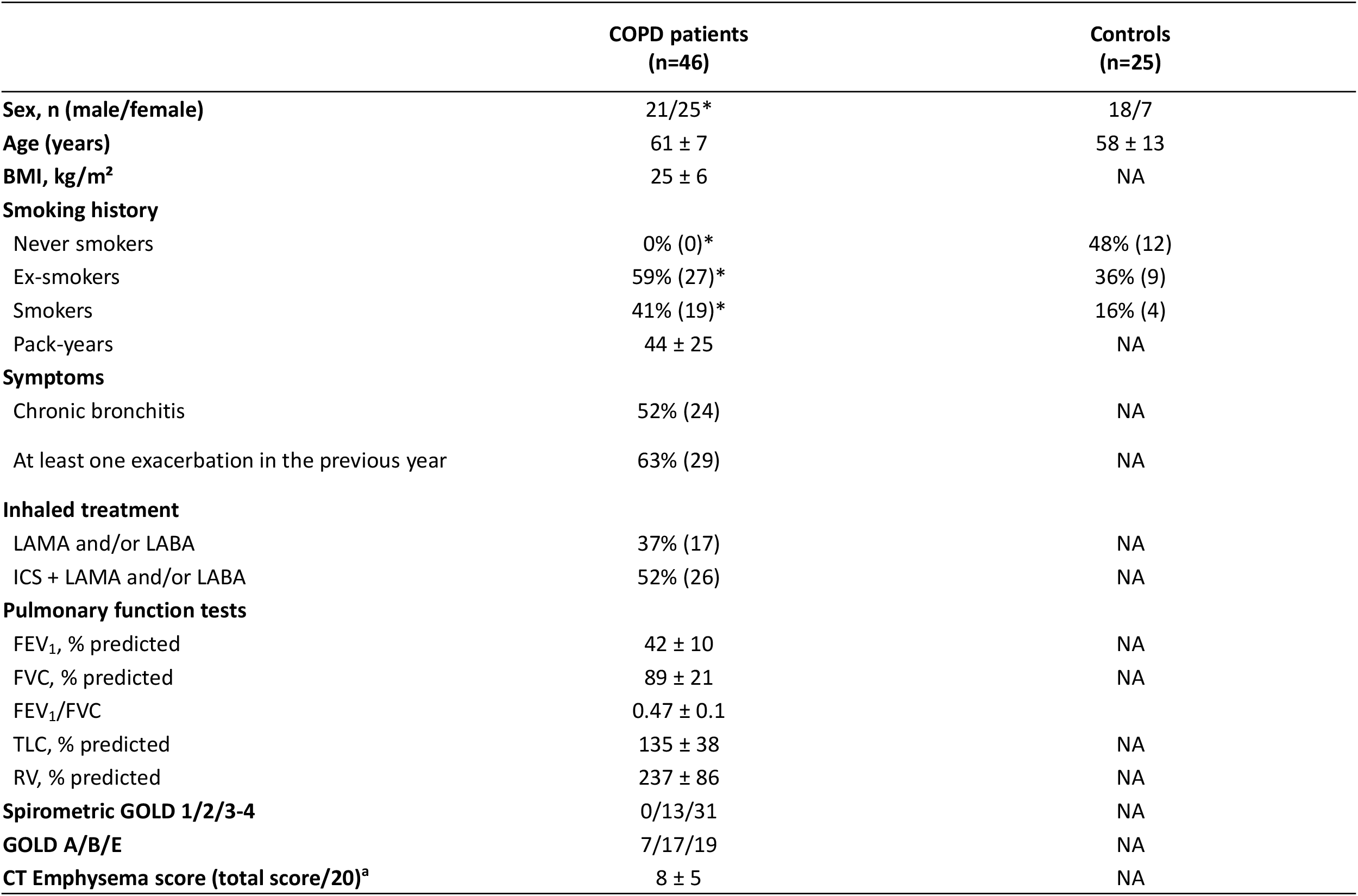
Characteristics of the study population. Values are expressed as mean ± Standard Error of the Mean (SEM) or percentage (number) **BMI** Body Mass Index, **CT** Computed tomography, **GOLD** Global Initiative for Chronic Obstructive Lung Disease, **LABA** long-acting beat-agonist, **LAMA** long-acting muscarinic antagonist, **NA** Not available, **ICS** inhaled corticosteroid, **FEV**_**1**_, forced expiratory volume in one second, **FVC** forced vital capacity, **TLC** total lung capacity, **RV** residual volume ^*^p < 0.05 vs control group ^a^Emphysema score was determined using a visual score assigned to each lobe, based on the extent of tissue destruction from 0 (no emphysema) to 4 (> 75% destruction)

|  | COPD patients<br>(n=46) | Controls<br>(n=25) |
| --- | --- | --- |
| Sex, n (male/female) | 21/25* | 18/7 |
| Age (years) | 61 ± 7 | 58 ± 13 |
| BMI, kg/m² | 25 ± 6 | NA |
| Smoking history |  |  |
| Never smokers | 0% (0)* | 48% (12) |
| Ex-smokers | 59% (27)* | 36% (9) |
| Smokers | 41% (19)* | 16% (4) |
| Pack-years | 44 ± 25 | NA |
| Symptoms |  |  |
| Chronic bronchitis | 52% (24) | NA |
| At least one exacerbation in the previous year | 63% (29) | NA |
| Inhaled treatment |  |  |
| LAMA and/or LABA | 37% (17) | NA |
| ICS + LAMA and/or LABA | 52% (26) | NA |
| Pulmonary function tests |  |  |
| FEV<sub>1</sub>, % predicted | 42 ± 10 | NA |
| FVC, % predicted | 89 ± 21 | NA |
| FEV<sub>1</sub>/FVC | 0.47 ± 0.1 |  |
| TLC, % predicted | 135 ± 38 | NA |
| RV, % predicted | 237 ± 86 | NA |
| Spirometric GOLD 1/2/3-4 | 0/13/31 | NA |
| GOLD A/B/E | 7/17/19 | NA |
| CT Emphysema score (total score/20)<sup>a</sup> | 8 ± 5 | NA |

### Samples collection and processing

Peripheral blood mononuclear cells (PBMC) from controls and COPD patients were collected from heparinized whole blood using a density gradient medium (Pancoll, PAN-Biotech). PBMC were washed with PBS (Biosera) two times before cytometry staining.Sera were harvested from the blood in dry tubes, retrieved after centrifugation for 10 minutes at 1200 g at 20°C and stored at −20°C until processing.

### Flow cytometry

Analysis of the surface and intracellular markers expression was performed using antibodies listed in Supplementary Table 1. For cytokine stainings, PBMC were stimulated for 2h with Phorbol myristate acetate (PMA) (Sigma, 1 µg/µL), Ionomycine (Sigma, 0,5 µg/µL) and Golgi Plug (BD Biosciences, 1 µg/µL). Surface staining was performed on 5×10^6^ cells for 30 minutes under agitation at 4°C in the dark. Cells were then treated with either transcription factors or intracellular cytokine staining kit (BD Biosciences) according to manufacturer’s instructions. Incubation with anti-cytokine and anti-transcription factor antibodies was respectively performed for 1 hour or overnight at 4°C under agitation in the dark. Cells were washed and resuspended in PBS. Flow cytometry was performed using BD LSRFortessa cell analyser (BD Biosciences). A minimum of 5000 ILC were acquired and analysed with Flowlogic v7.3 software (Inivai Technologies).

### PBMC-derived macrophages culture and stimulation

Monocytes from Pancoll-isolated PBMCs were differentiated into macrophages (MDMs) by culturing in X-VIVO 15 medium (Lonza) with 10% foetal bovine serum (Gibco), 100 U of penicillin/mL, 100 µg of streptomycin/mL (Gibco) and 50 ng/mL of M-CSF (Miltenyi Biotec) for 5 days. MDMs were stimulated with scrambled peptide (SP) (10 µg/mL, Genepep) or elastin peptide VGVAPG (10 µg/mL, GenScript) for 24 to 48h before collecting the supernatant. 400 µL of TRIzol (Invitrogen) were added to each well and macrophages lysates were collected.

### ILC enrichment and culture

ILC were isolated from PBMCs by negative selection using the EasySep™ Human Pan-ILC Enrichment Kit according to manufacturer’s instructions (Stemcell Technologies). ILC were sorted using FACSARIA™IIu (BD Biosciences) and cultured in 96-well culture plates (2×10^3^ cells/well) with serum or EP-stimulated MDMs supernatants in X-VIVO 15 medium (Lonza) with 10 U/mL of IL-2 (PeproTech) and 50 ng/mL of IL-7 (PeproTech).

### Single cell RNA-seq processing

Publicly available scRNA-seq datasets [16, 17] comparing lung tissue from patients with or without COPD were analysed. Dataset were processed using Cellenics® (https://scp.biomage.net/) with the parameters described in Supplementary table 2.

### RT-qPCR

RNA from MDMs was isolated according to the manufacturer’s protocol. cDNA was prepared using Maxima First Strand cDNA Synthesis Kit (Thermofisher Scientific) according to the manufacturer’s protocol. RT-PCR reactions were performed in a 20 μL mixture containing 1× Power SYBR™ Green PCR Master Mix (Thermofisher Scientific), 0.5 μmol/L of each primer, and 4 μL of cDNA template. Primers (Thermofisher Scientific) used for detection of human *IL4, IL5, IL13, IL25, IL33, AREG*, and *GAPDH* are listed in Supplementary Table 3. Real-time PCR was performed using the Applied Biosystem 7500 Fast system under the following cycling conditions: 95 °C for 10 min, 40 cycles of 95 °C for 15 s, and 60 °C for 1 min, followed by the melting curve stage. The relative mRNA expression level was normalized to *GAPDH* using the 2^-ΔΔCT^ method.

### Quantification of cytokines

Levels of IL-4, IL-5 and IL-13 were measured in control subject and COPD patients’ serum and SP or EP-stimulated MDMs supernatants using LEGENDplex HU Th Panel according to manufacturer’s instructions (BioLegend). Aquisition was performed on Accuri C6 Flow Cytometer (BD Biosciences) and analyzed by the LEGENDplex Software.

### Statistics

Data are presented as mean ± SEM. Data analysis was performed using Prism 9 (GraphPad). Comparisons were performed either with Student’s t-test, Wilcoxon-Mann–Whitney test one-way ANOVA with Tukey’s multiple comparison test. P values < 0.05 were considered significant with *<0.05, **<0.01 and ***<0.001.

## RESULTS

### COPD patients have an type2-ILC profile

We conducted ILC analyses based on the gating strategy presented in Supplementary figure 1. Clinical data of both the COPD patients and the control subjects were analyzed (Table 1). Differences were found in the sex distribution and, due to the well-established association between COPD and cigarette smoking, smoking status between COPD patients and controls. However, no significant differences were found between never-smokers and ever-smokers in the control group (Supplementary figure 2), and the analyses of sex differences and correlations with emphysema scores did not reveal any statistically significant differences (Supplementary figures 3 and 4).

We observed a comparable proportion of total ILC and ILC1 between COPD patients and control subjects (Figure 1A and 1B). Conversely, there was a significant increase in the proportion of ILC2, and a decrease in the proportion of ILC3 in the peripheral blood of COPD patients (Figure 1B). Moreover, our analysis unveiled an elevation in the proportion of IFN-y-expressing ILC1, IL-4 expressing ILC2, and IL-17-expressing ILC3 in COPD patients compared to controls (Figure 1C). We then examined patients’ clinical data and showed that the total proportion of ILC and the proportion of IL-4-secreting ILC2 were diminished in smoking COPD patients compared to ex-smokers (Figure 1D and E). Additionally, we showed a significant decrease in the proportion of IL-4-secreting ILC2 specifically in COPD patients with chronic bronchitis (Figure 1F).

**Figure 1:**
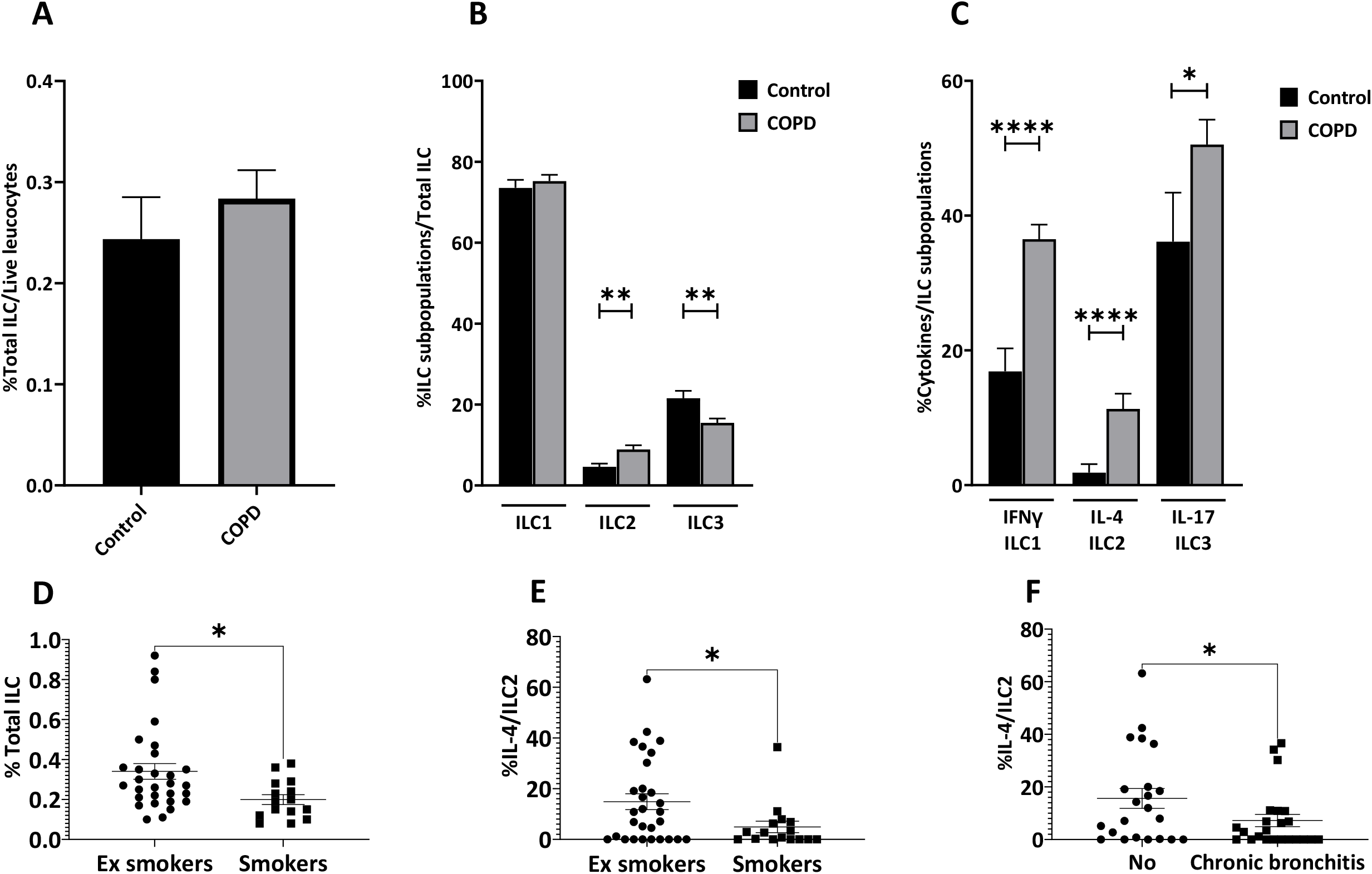
COPD patients have an ILC2 profile. **(A)** Proportion of total ILC. **(B)** Proportion of ILC subpopulations. **(C)** Proportion of ILC1, ILC2 and ILC3 subpopulations secreting IFN-γ, IL-4 and IL-17 respectively. (D-E) Proportion of total ILC **(D)** and of IL-4 secreting ILC2 **(E)** regarding smoking history of COPD patients. **(F)** Proportion of IL-4 secreting ILC2 in COPD patients suffering or not from chronic bronchitis. ILC isolated from the blood of COPD patients (n=46) and controls (n=25) were used for phenotypic and functional characterization by flow cytometry. Data are presented as mean ± SEM. Differences between two groups were evaluated by using the Wilcoxon-Mann–Whitney test unless otherwise specified. P values <0.05 were considered significant.

### COPD sera orientate ILC from control subjects toward a COPD-like profile

To further assess the influence of COPD on ILC, we enriched ILC from controls and cultured them with either IL-2 and IL-7 (as a control of baseline survival), 10% control serum or 10% COPD serum. We showed that the proportion of ILC1 and ILC3 remained unaffected after 5 days of culture with 10% COPD serum in comparison to control serum (Supplementary figure 5A and 5C). We observed a significant increase in the proportion of ILC2 after incubation with 10% COPD serum (Supplementary figure 5B). We then assessed the presence of specific factors that may favour ILC2 in patients’ *sera*. Cytometry-sorted ILC were cultured with IL-2 and IL-7, to give a consistant survival signal, and in the presence or absence of 10% COPD serum. Remarkably, the proportions of ILC1 and ILC2 were significantly increased after 5 days of culture in the presence of 10% COPD serum (Figure 2A and 2B), while the proportion of ILC3 was decreased (Figure 2C).

**Figure 2:**
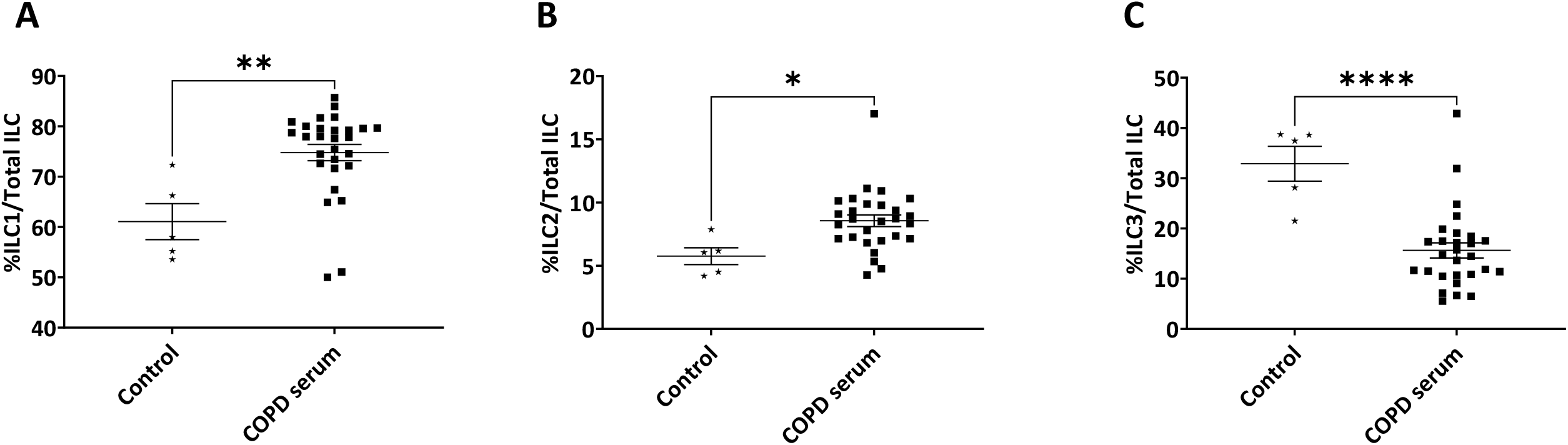
COPD serum can orientate healthy ILC toward a COPD profile. **(A)** Proportion of ILC1. **(B)** Proportion of ILC2. **(C)** Proportion of ILC3. Sorted ILC isolated from the blood of controls (n=5) were cultured for 5 days in the presence of IL-2 and IL-7 (control) or IL-2 + IL-7 and 10% COPD serum isolated from patients (n=27) before being analyzed by flow cytometry. Data are presented as Mean±SEM. Differences between two groups were evaluated by using the Wilcoxon-Mann–Whitney test. P values <0.05 were considered significant.

### EP-stimulated macrophages favor ILC2

We hypothetize that the extensive degradation of lung elastin in COPD patients, resulting in high concentrations of EP in the lungs and blood of COPD patients, might have an impact on ILC. We determined that ILC1, ILC2 and ILC3 do not express EP receptors (Figure 3A) in blood ILC from neither controls nor COPD patients (Figure 3B). Consequently, we decided to investigate which cell type express EP receptors and could potentially impact ILC. To achieve this, we analysed publicly available scRNAseq datasets from COPD patients and controls [16, 17] (Figure 3C). We examined EP receptors genes: *CTSA, NEU1*, and *GLB1*, encoding the Elastin Receptor Complex, *LGALS3* which codes for galectin-3, and integrins alpha V (*ITGAV*), beta 3 (I*TGB3*), and beta 5 (*ITGB5*). We showed that among immune cells, genes encoding the ERC and galectin-3 were mostly expressed by alveolar and interstitial macrophages (Figure 3D) but not differentially expressed between COPD patients and controls (Figure 3E and 3F).

**Figure 3:**
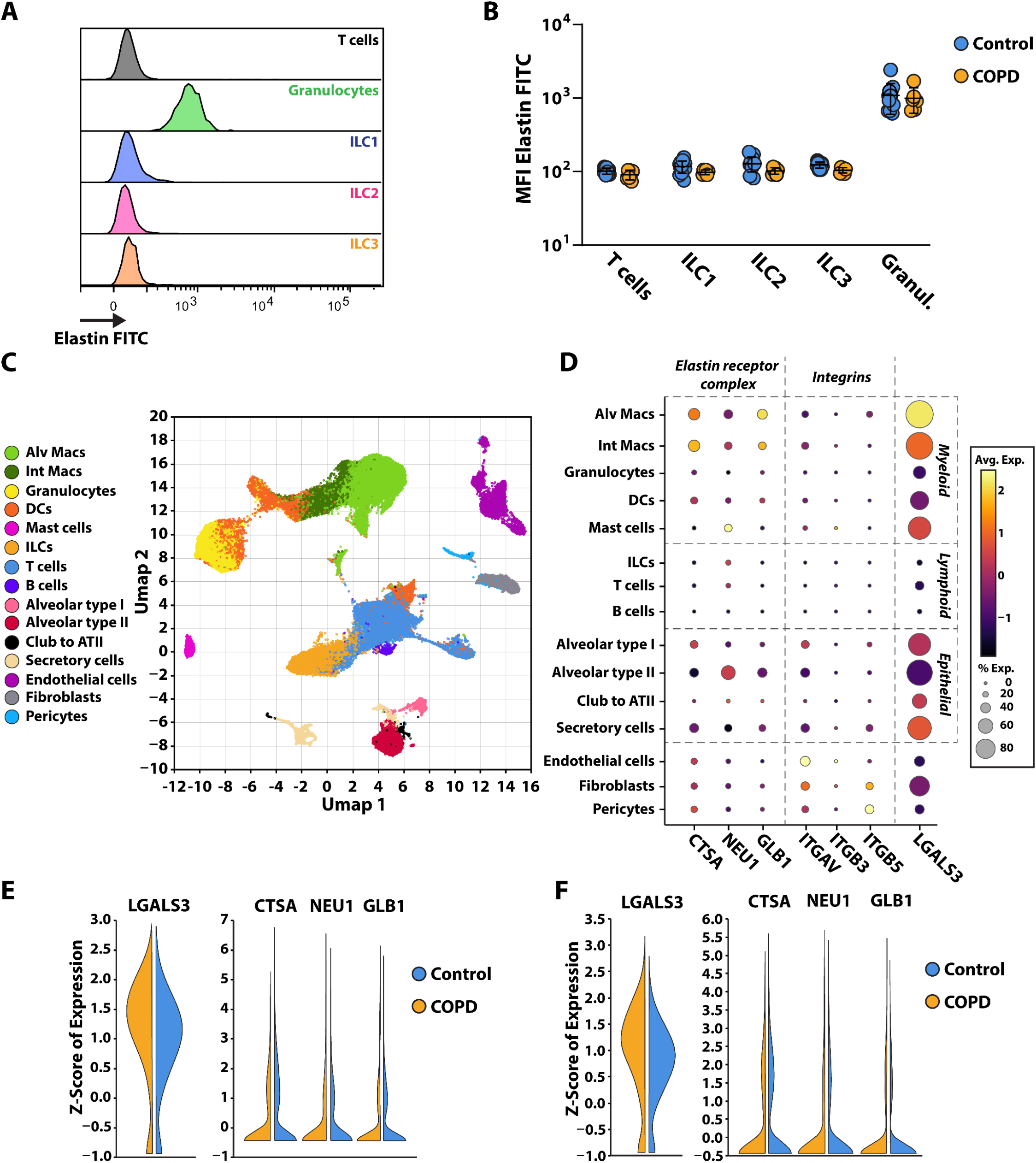
Lung macrophages express EP receptors but not ILC. **(A)** Overlays of immune cells expression of elastin receptor. **(B)** Comparison of FITC-conjugated elastin MFI in immune cells of COPD patients (n=6) and control subjects (n=12). **(C)** UMAP showing the 15 clusters obtained with GO reports and signaling pathways in the lungs. **(D)** Representation of the expression of genes associated with elastin by the different clusters of the lungs. **(E-F)** Expression of EP receptors on alveolar **(E)** and interstitial **(F)** macrophages of COPD patients and control subjects.

Therefore, we examined the role of EP-stimulated macrophages on ILC. We cultures ILC for 5 days with supernatants coming from SP or EP-stimulated monocyte-derived macrophages Our results revealed that neither the supernatant from 24h nor the supernatant from 48h EP-stimulated macrophages influenced the proportion of ILC1 or ILC3 (Figure 4A and 4C) while the proportion of ILC2 was increased with 48h EP-stimulated macrophage supernatants culture (Figure 4B). This suggest that EP modulate ILC through their impact on macrophages.

**Figure 4:**
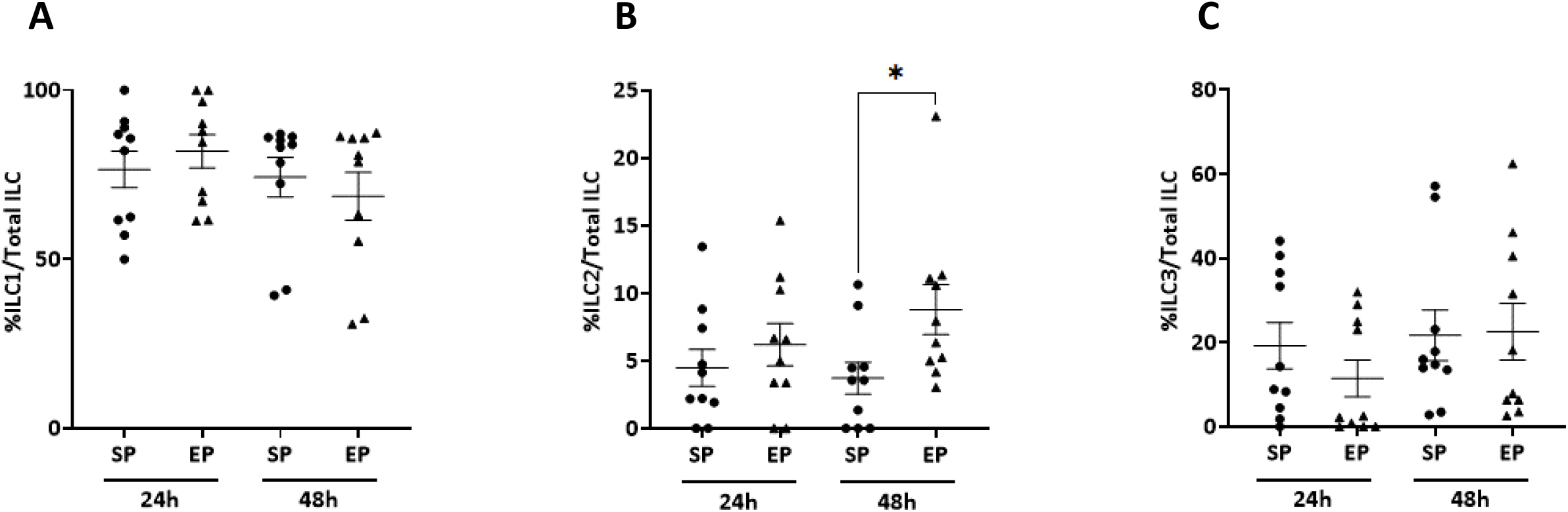
EP-stimulated macrophages modulate ILC2 expansion. **(A-C)** Sorted ILC isolated from the blood of control subjects (n=2) were cultured with scrambled peptide (SP) or elastin peptide (EP) stimulated macrophage supernatant (n=10) before being analyzed by flow cytometry for ILC1 (A), ILC2 (B) and ILC3 (C) proportion. Sorted ILC isolated from the blood of controls (n=2) were cultured for 5 days with scrambled peptide or elastin peptide stimulated macrophages’ supernatant (n=10) before being analyzed by flow cytometry. Data are presented as Mean±SEM. Differences between two groups were evaluated by using the Wilcoxon-Mann–Whitney test. P values <0.05 were considered significant.

### ILC2 frequency is correlated with IL-5 secretion by EP-stimulated macrophages and COPD patients’ *sera*

We next analysed the mRNA expressions of cytokines known to influence ILC2 orientation. The expression of *IL25, IL33*, and *AREG* were not detected but we observed a significant increase in the expression of *IL4* (Figure 5A), *IL5* (Figure 5B), and *IL13* (Figure 5C) in EP-stimulated macrophages compared to SP. We analysed IL-4, IL-5, and IL-13 in the culture supernatants. EP stimulation did not affect the secretion of IL-4 (Figure 5D), IL-13 was not detectable (Figure 5F) and only IL-5 was increased (Figure 5E). We then assessed the concentrations of IL-4, IL-5, and IL-13 in controls’ and COPD patients’ *sera*. We demonstrated a significant elevation in the levels of IL-5 and IL-13 in the *sera* of COPD patients (Figure 6A-C). However, a correlation between ILC2 frenquency and cytokine concentration was only found for IL-5 (Figure 6D-F) supporting its pivotal role in the promotion of ILC2.

**Figure 5:**
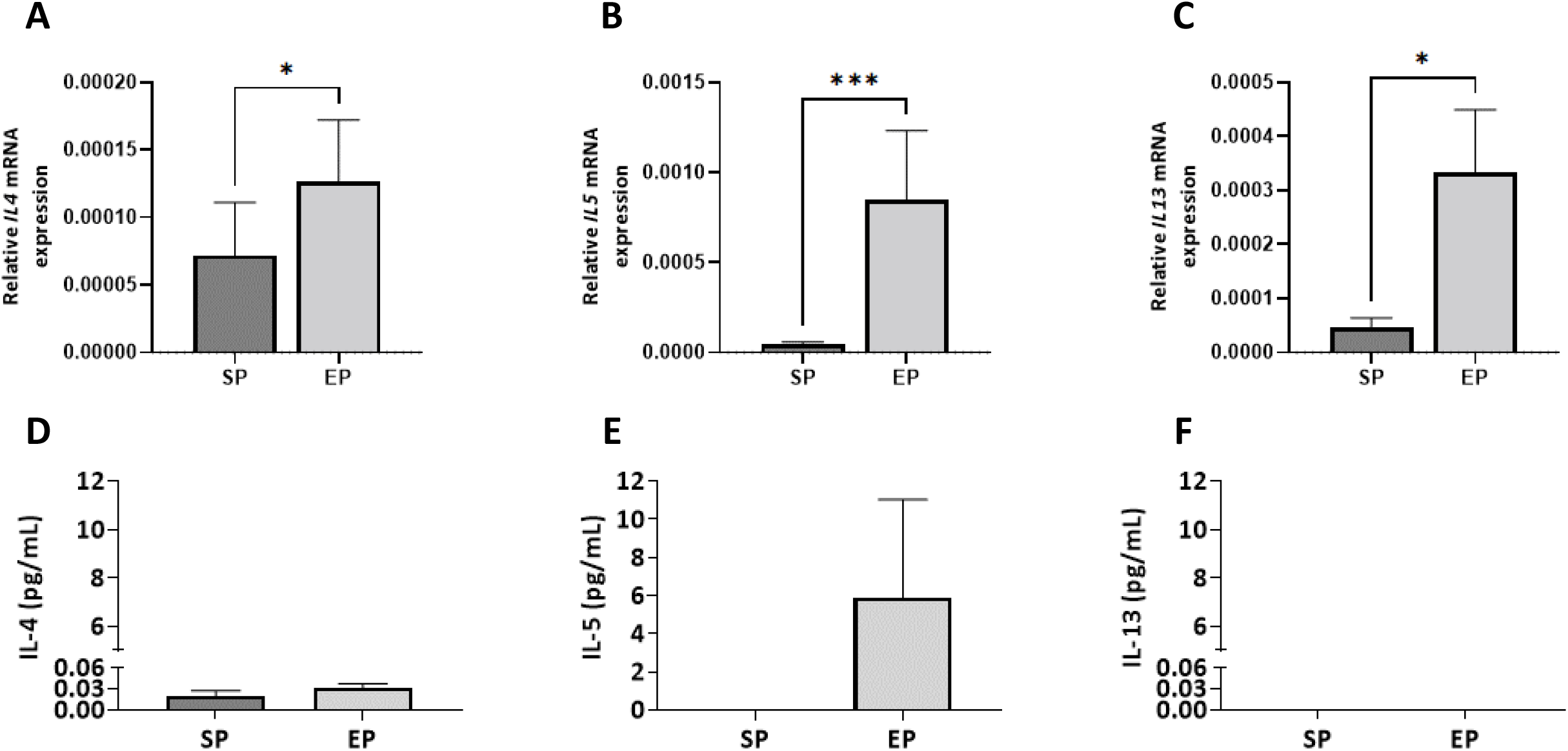
ILC2 expansion is correlated with IL-5 secretion by EP-stimulated macrophages. **(A-C)** Relative mRNA expression of IL-4 **(A)**, IL-5 **(B)** and IL-13 **(C)** in 24h SP- or EP-stimulated macrophages. **(D-F)** IL-4 **(D)**, IL-**5 (E)** and IL-13 **(F)** concentrations in 48h stimulated macrophage supernatants. PBMC were isolated from the blood of patients and controls (total n=11) and monocytes were differentiated into macrophages for 5 days by culturing them with M-CSF. SP or EP was added to the media for 24 or 48h. Macrophages RNA were extracted and RTqPCR was performed. Cytokines quantification was performed using LEGENDplex HU Th Panel on 48h SP- or EP-stimulated macrophages supernatants (n=4-6/group). Data are presented as Mean±SEM. Differences between two groups were evaluated by using the Wilcoxon-Mann–Whitney test. P values <0.05 were considered significant.

**Figure 6:**
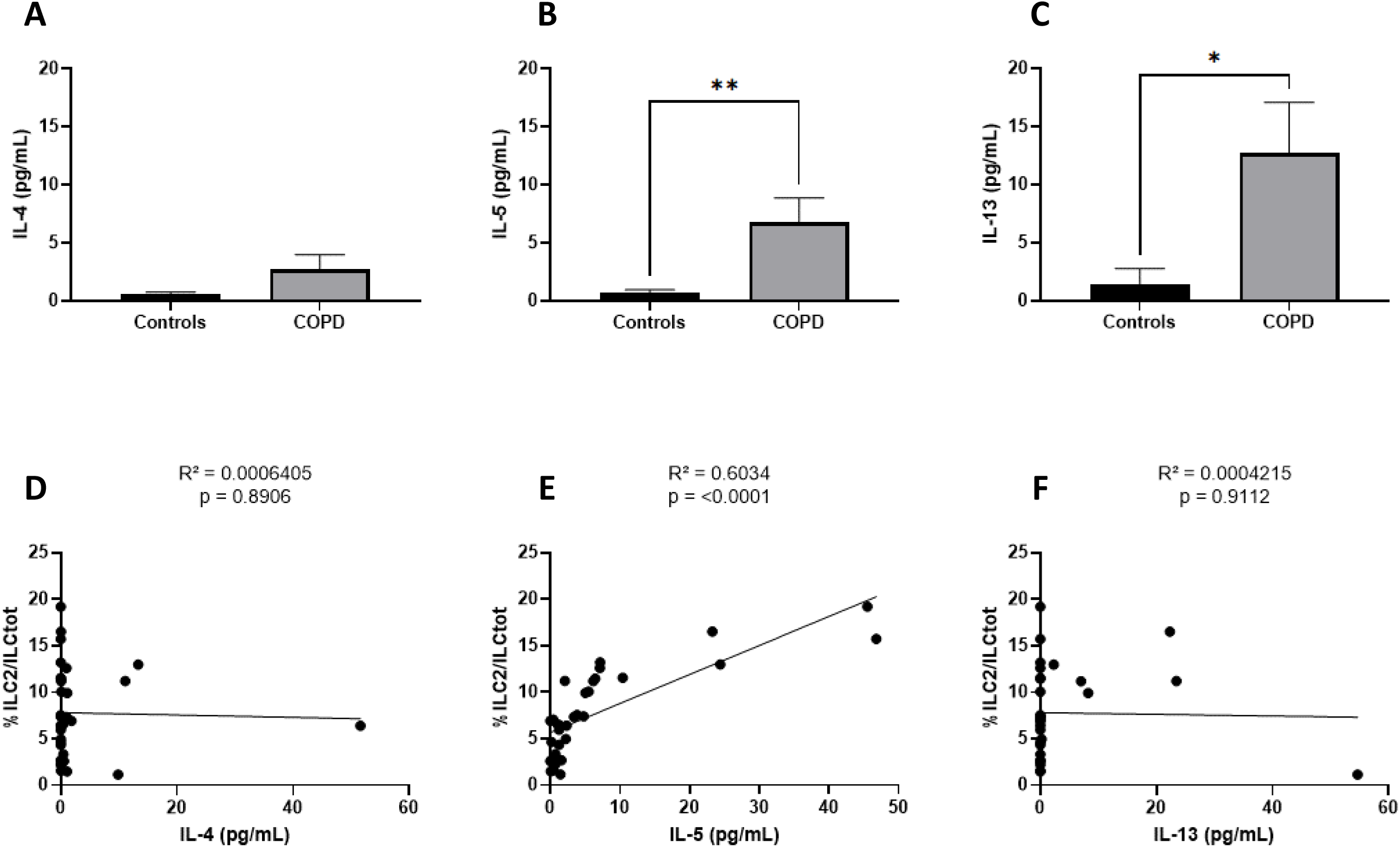
IL-5 in COPD serum correlate with ILC2 expansion. **(A-C)** IL-4 **(A)**, IL-5 **(B)** and IL-13 **(C)** concentration in sera from control subjects (n=13) and COPD patients (n=32). **(D-F)** Pearson correlation between ILC2 proportion and IL-4 **(D)**, IL-5 **(E)** and IL-13 **(F)** concentration in COPD patients. Cytokines quantification was performed using LEGENDplex HU Th Panel on controls (n=13) and COPD patients (n=32) sera. Data are presented as Mean±SEM. Difference between two groups were evaluated by using the Wilcoxon-Mann–Whitney test. Pearson coefficient was used for correlation analysis. P values <0.05 were considered significant.

## DISCUSSION AND FUTURE DIRECTIONS

In this study, we performed the phenotypic and functional analysis of ILC in COPD patients. We showed an increase in the proportion of ILC2, a decrease in ILC3 and a general activation of ILC in COPD patients compared to controls. The use of COPD sera resulted in an increase in the proportion of ILC2. Because elastin breakdown in the lung of COPD patients leads to the release of high concentrations of EP that elicit a biological impact on immune cells [4, 6], we here hypothesized that EP could also have an impact on ILC. However, we found EP receptors expressed on lung macrophages but not on ILC. Furthermore, we suggest an indirect role of EP on ILC2 through their effect on IL-5 secretion by macrophages. Finally, the concentration of IL-5 was elevated in the *sera* of COPD patients and correlated with the proportion of ILC2. These results highlight the role of EP, macrophages, and IL-5 in modulating ILC responses in the context of COPD.

Our phenotypic and functional studies demonstrate a shift of ILC towards an ILC2 profile in the peripheral blood of COPD patients, consistent with previous research [14]. We further observed a general activation of ILC indicated by an increased capacity of cytokine secretion. An elevated proportion of ILC1 secreting IFN-y, which has been linked to emphysema development in animal models [18], of ILC2 secreting IL-4, known for its role in tissue repair and homeostasis [19], and of ILC3 secreting IL-17, involved in neutrophil maturation and recruitment [20], was found in peripheral blood from COPD patients. Furthermore, these cytokines have been shown to be implicated in promoting mucus secretion and alveolar damage through stimulating the release of proteases, underscoring their importance in the pathophysiology of COPD [18–20].

Interestingly, we suggest that cigarette smoking may exert an influence on the profile of innate lymphoid cells in individuals with COPD. Specifically, we observed a decrease in both the total proportion of ILC, as previously reported in a study on asthma [21], and the proportion of ILC2 secreting IL-4 in smokers compared to non-smokers with COPD and patients with chronic bronchitis. Previous research has indicated that smoking can reduce the constitutive levels of genes associated with Th2 cells, such as *Gata3, Il4ra*, and *Postn* in the lungs [22] and have linked active cigarette smoking and chronic bronchitis [23]. These findings highlight the need for further investigations of the role of ILC2 in the physiopathology of COPD and the impact of smoking on ILC profiles in the context of COPD. While ILC2 are clearly pathogenic in allergic asthma [24], their exact role in COPD remains unclear probably due to the mixed type 1/type 2 nature of the inflammatory response found in COPD. Indeed, on the one hand it has been shown that ILC2 can participate to COPD pathogenesis by promoting type 2 inflammation during exacerbations [25] and by converting to ILC1 that promote emphysema [26]. On the other hand, it has been shown in a cigarette-smoke induced COPD model that the absence of ILC2 favors lung fibrosis [15] and can potently secrete IL-10, adopting an ILC regulatory phenotype both in human and mice [27–29].

Furthermore, our study reveals intriguing evidence that COPD serum possesses the potential to induce a shift in the profile of innate lymphoid cells from healthy subjects towards a COPD-associated phenotype. Specifically, we observed a significant increase in the proportion of ILC2 after culture with COPD serum, which aligns with a previous study by *Chen et al*. [30] suggesting the influence of serum factors on ILC profiles. These findings collectively strengthen the notion that specific factors present in the serum of COPD patients have the capacity to modulate the behaviour and phenotype of ILC.

In individuals with COPD, there is a significant breakdown of lung elastin following the destruction of the alveoli, leading to increased concentrations of EP locally and systemically. To gain deeper insights into the intricate relationships between ILC and EP, we analysed publicly available scRNAseq datasets. We identified macrophages as potential key effectors due to their high expression of EP receptors. Notably, during emphysema, resident pulmonary macrophages exhibit a propensity to aggregate in emphysematous alveolar regions and initiate early responses [31]. This observation suggest a potential role of macrophages in in orchestrating the intricate interplay between EP and ILC. To further explore this connection, we cultured ILC with supernatant from macrophages that had been stimulated with EP. We observed a significant increase in the proportion of ILC2 suggesting that macrophages may play a regulatory role on ILC2 cells through cytokines secretion. Moreover, macrophages and ILC2 cells are known to play critical roles in various lung diseases, including parasitic infections, asthma, and cancers. In fact, it has been shown that M2 macrophages promote ILC2 activation and differentiation from progenitors in asthma [32, 33]. Our findings highlight further the importance of studying the macrophages and ILC2 crosstalk in pulmonary diseases, especially in COPD.

To identify the factor(s) responsible for promoting ILC2 under EP stimulation, we investigated the expression of cytokines known to favor them. We found that ILC2 promotion was correlated with IL-5 secretion by EP-stimulated macrophages and patients IL-5 *sera* concentrations. IL-5 is well known to be a major mediator of eosinophilic inflammation during lung conditions especially in asthma where it plays a major pathogenic role [34]. In COPD, the role of IL-5 and its potential targeting remains a conundrum. Indeed, some studies advocate for the use of anti-IL-5/IL-5R treatment [35–37] based on benefits on exacerbations while others call for more research to identify patients that would benefit from IL-5 targeting [38–40].

This study provides new elements in understanding of COPD pathophysiology. It establishes the critical interplay between elastin peptides and ILC2 in COPD. These insights pave the way for potential therapeutic interventions targeting this pathway in COPD management. This study is limited by our focus on stable COPD patients. Also, we did not explore the predictability of IL-5, EP concentrations or ILC2 rate on patients’ clinical follow-up. Further investigation is warranted to comprehensively address the role of EP during COPD exacerbations.

## Notes

### Competing Interest Statement

The authors have declared no competing interest.

