## Supplementary material for "Elastin-derived peptides favor type 2 innate lymphoid cells in COPD"

| Antibody | Clone | Fluorochrome | Reference and manufacturer |
|---|---|---|---|
| Lineage | UCHT1; HCD14; 3G8; HIB19; 2H7; HCD56 | Pacific Blue | Biolegend - 348805 |
| CD7 | M-T701 | PE CF594 | BD Biosciences – 562541 |
| CD127 | HIL-7R-M21 | PE CY7 | BD Biosciences – 560822 |
| CD294/CRTH2 | BM16 | ALEXA FLUOR 647(APC) | BD Biosciences – 558042 |
| CD117 | 104D2 | BV605 | BD Biosciences – 562687 |
| ElastinR | HP59 | FITC | Elastin Products Company |
| Zombie NIR | | APC/Cy7 | Biolegend - 423105 |
| Tbet | AD2 | PE | BD Biosciences - 561258 |
| GATA3 | L50-823 | BB700 | BD Biosciences - 566642 |
| RORyt | REA278 | Vio-B515 | Miltenyi Biotec - 130-123-840 |
| IL-4 | MP4-25D2 | APC-Cy7 | Biolegend - 500834 |
| IL-17 | BL168 | PerCP/Cy5.5 | Biolegend - 512314 |
| IFNy | B27 | PE | BD Biosciences - 559327 |
| CD3 | UCHT1 | BV421 | BD Biosciences – 562426 |
| CD4 | SK3 | PECy7 | BD Biosciences – 348809 |
| CD8 | SK1 | BV510 | BD Biosciences - 563919 |

**Supplementary Table 1:** Flow cytometry antibodies

| Step number | Step | Parameters and values |
| --- | --- | --- |
| 1 | Classifier | FDR=0.01 |
| 2 | Cell Size Distribution | binStep=200, minCellSize=500 |
| 3 | Mitochondrial Content | Method=absoluteThreshold, binStep=0.3, maxFraction=0.1 |
| 4 | NumGenes Vs NumUmis | RegressionType=linear, p.level=0.0002221235 |
| 5 | Doublet Scores | binStep=0.02, ProbabilityThreshold=0.2 |
| 6 | Data Integration | Method=harmony, NumGenes=2000, Normalisation=logNormalize |
| 7 | Dimensionality Reduction | Method=rpca, numPCs=30, ExcludeGeneCategories=cellCycle |
| 8 | Embedding Settings | Method=umap, distanceMetric=cosine, minimumDistanc=0.3 |
| 9 | Clustering Settings | Method=Louvain, resolution=1.2 |

**Supplementary Table 2:** Pipeline used for scRNAseq analysis

| Primer | Sequence |
|---|---|
| *IL4* F | CCGTAACAGACATCTTTGCTGCC |
| *IL4* R | GAGTGTCCTTCTCATGGTGGCT |
| *IL5* F | GGAATAGGCACACTGGAGAGTC |
| *IL5* R | CTCTCCGTCTTTCTTCTCCACAC |
| *IL13* F | ACGGTCATTGCTCTCACTTGCC |
| *IL13* R | CTGTCAGGTTGATGCTCCATACC |
| *IL25* F | AACCGCCACCCAGAGTCCTGT |
| *IL25* R | ACAGGCAACGGGCGTGGTACA |
| *IL33* F | GCCTGTCAACAGCAGTCTACTG |
| *IL33* R | TGTGCTTAGAGAAGCAAGATACTC |
| *AREG* F | GCACCTGGAAGCAGTAACATGC |
| *AREG* R | GGCAGCTATGGCTGCTAATGCA |
| *GAPDH* F | GTCTCCTCTGACTTCAACAGCG |
| *GAPDH* R | ACCACCCTGTTGCTGTAGCCAA |

**Supplementary Table 3:** RT-qPCR primers

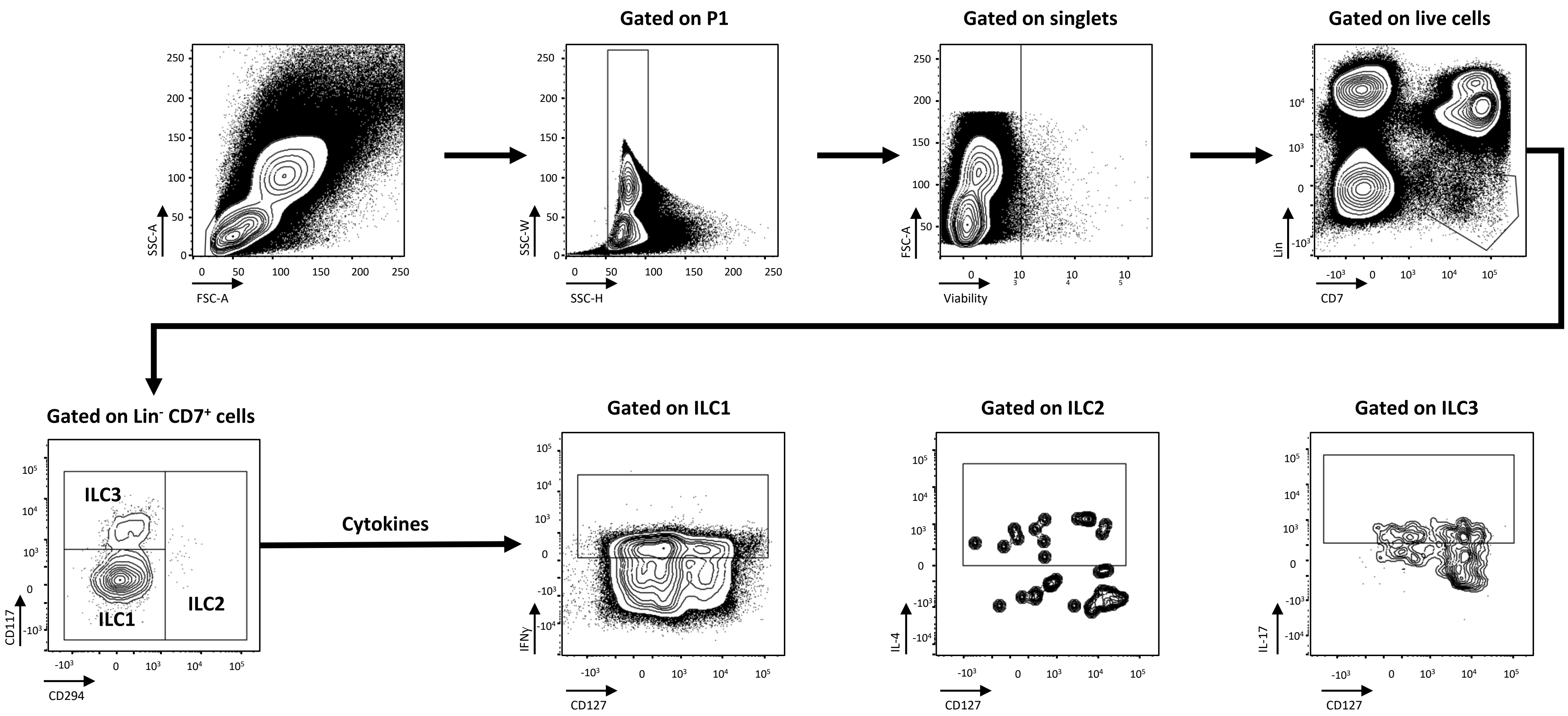


**Supplementary figure 1:** Gating strategy used for cytometry analysis.

The size and complexity of peripheral blood mononuclear cells were used to select leukocytes. After exclusion of doublets and dead cells, innate lymphoid cells were defined as $Lin^-$ and $CD7^+$. Lineage (CD3/14/16/19/20/56) allowing us to eliminate T and B lymphocytes, natural killer cells and myeloid cells. Among ILC, the expression of the CD117 and CD294 markers was used to determine the 3 sub-populations: ILC1 ($CD117^-$ and $CD294^+$), 2 ($CD117^{+/-}$ and $CD294^+$) and 3 ($CD117^+$ and $CD294^-$). The expression of their associated secreting cytokines were also analysed. i.e., IFN-$\gamma$ for ILC1, IL-4 for ILC2 and IL-17 for ILC3.

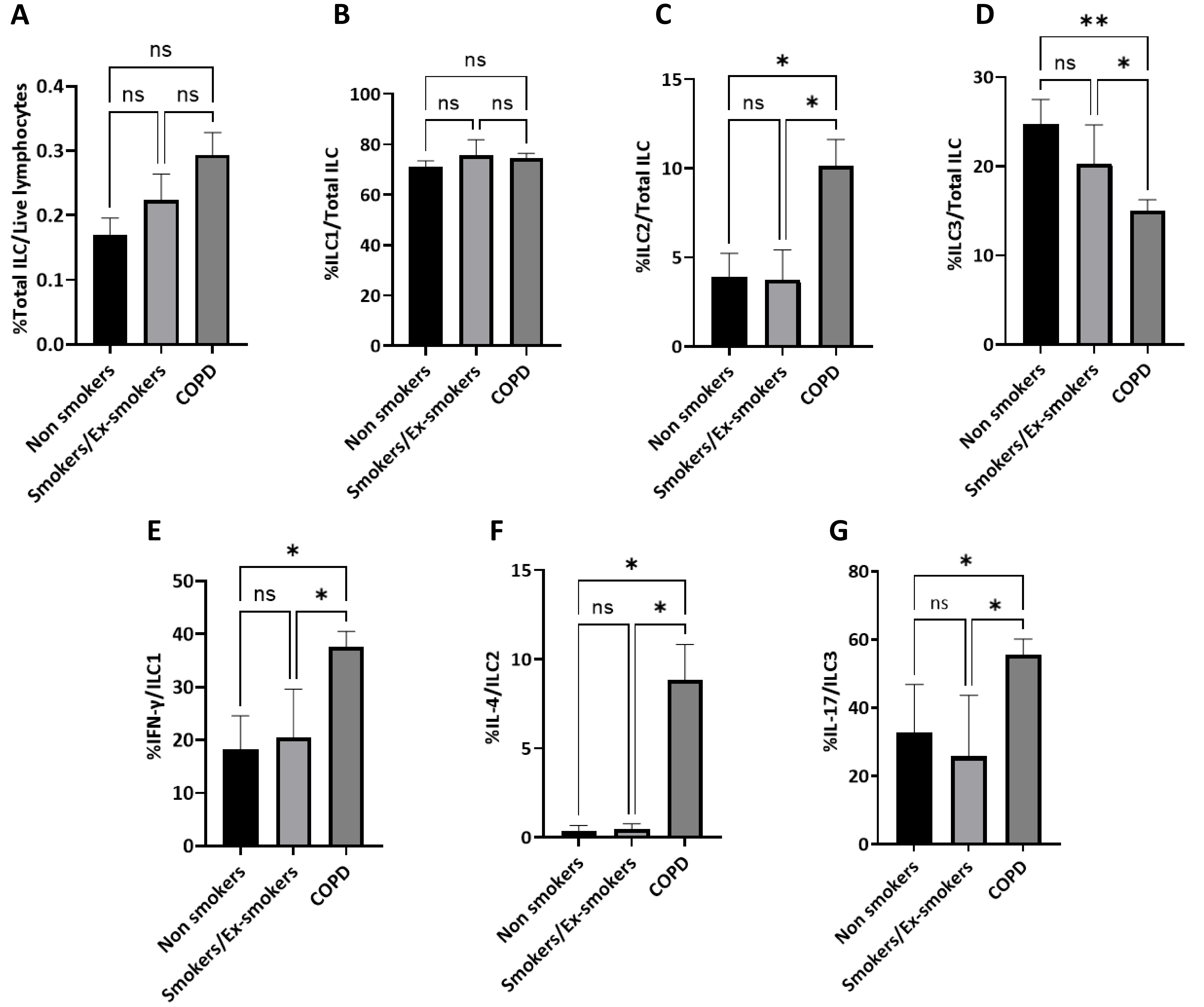


**Supplementary figure 2:** Comparison of **(A)** Total ILC proportion, **(B)** ILC1 proportion, **(C)** ILC2 proportion, **(D)** ILC3 proportion, **(E)** Proportion of IFN-γ secreting-ILC1, **(F)** Proportion of IL-4 secreting-ILC2, **(G)** Proportion of IL-17 secreting-ILC3 between non-smokers (n=12), ex-smokers and current smokers' controls (=13) and COPD patients. Data are presented as mean ± SEM. Differences between two groups were evaluated by using the Wilcoxon-Mann–Whitney test unless otherwise specified. P values <0.05 were considered significant.

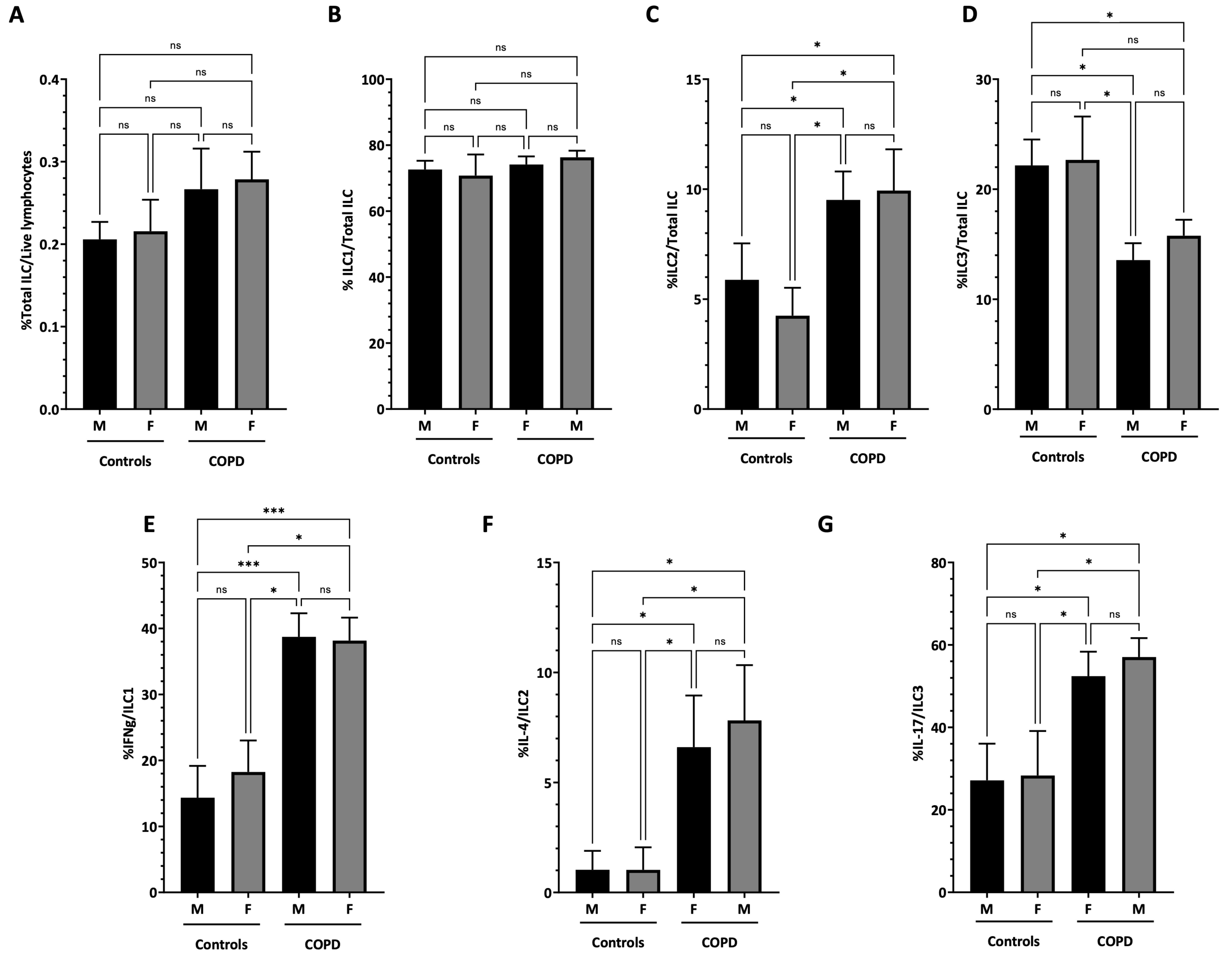


**Supplementary figure 3:** Comparison of **(A)** Total ILC proportion**, (B)** ILC1 proportion**, (C)** ILC2 proportion, **(D)** ILC3 proportion**, (E)** Proportion of IFN-γ secreting-ILC1, **(F)** Proportion of IL-4 secreting-ILC2, **(G)** Proportion of IL-17 secreting-ILC3 between male (M) (n=18) and female (F) (n=17) controls and male (n=21) and female (n=25) COPD patients. Data are presented as mean ± SEM. Differences between two groups were evaluated by using the Wilcoxon-Mann–Whitney test unless otherwise specified. P values <0.05 were considered significant.

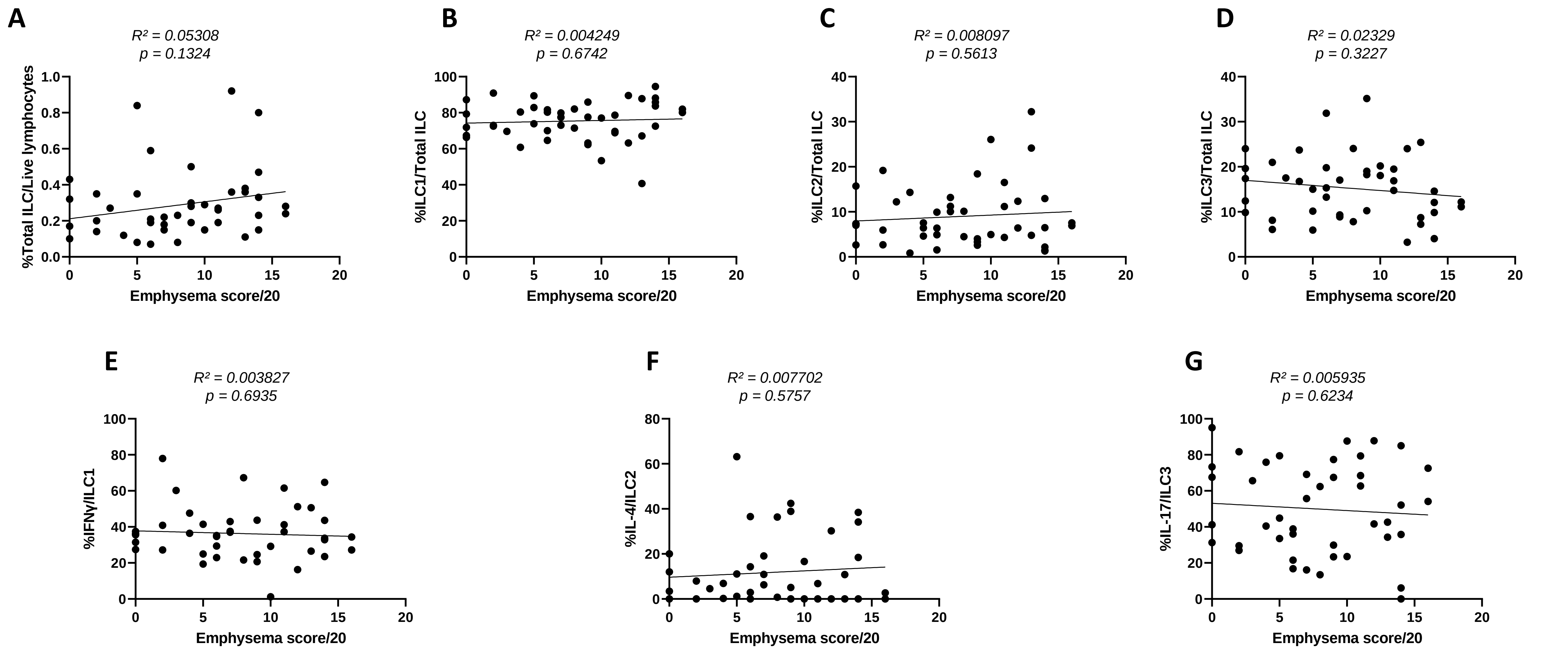


**Supplementary figure 4:** Pearson correlation between emphysema score out of 20 and proportion of total ILC **(A)**, ILC1 **(B)**, ILC2 **(C)**, ILC3 **(D)** and ILC1, ILC2 and ILC3 subpopulations secreting IFN-γ, IL-4 and IL-17 respectively **(E-G)** in COPD patients. Pearson coefficient was used for correlation analysis of COPD patients (n=46). P values <0.05 were considered significant.

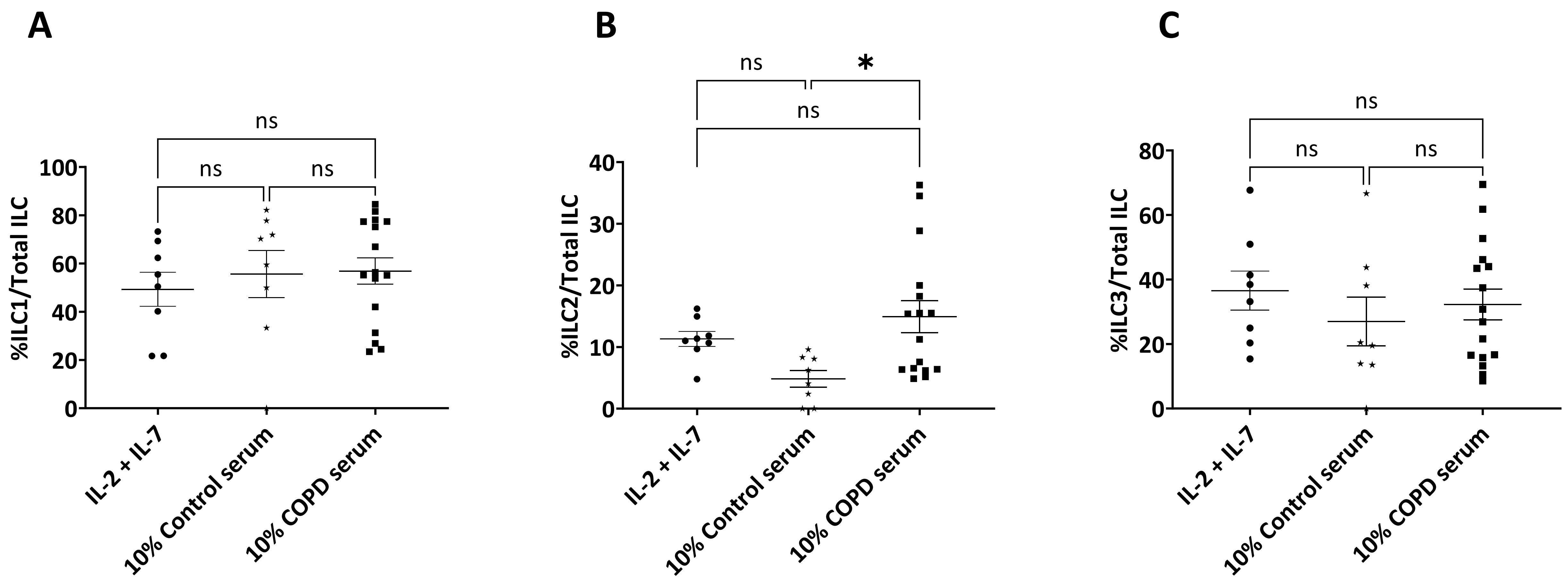


**Supplementary figure 5: (A)** Enriched proportion of ILC1. **(B)** Enriched proportion of ILC2**. (C)** Enriched proportion of ILC3. Enriched ILC isolated from the blood of controls (n=2-8) were cultured for 5 days in the presence of either IL-2 and IL-7 (n=8), 10% control serum (n=8) or 10% COPD serum isolated from patients (n=27) before being analyzed by flow cytometry. Data are presented as Mean±SEM. Significance between three groups was calculated using one-way ANOVA with Tukey's multiple comparison test. P values <0.05 were considered significant.
